# Radicular and periodontal structural defects underlie refractory oral pathology in the adult *Hyp* mouse model of X-linked Hypophosphatemia

**DOI:** 10.64898/2026.04.27.719778

**Authors:** Chiaki Nishizawa, Jiro Miura, Tomoaki Iwayama, Miwa Yamazaki, Toshimi Michigami, Kazuaki Miyagawa

**Affiliations:** Department of Bone and Mineral Research, Osaka Women’s and Children’s Hospital, Izumi, Japan; Division for Interdisciplinary Dentistry, Graduate School of Dentistry, Osaka University, Suita, Japan; Department of Periodontology and Regenerative Dentistry, Graduate School of Dentistry, Osaka University, Suita, Japan

**Keywords:** X-linked Hypophosphatemia (XLH), *Hyp* mouse, periodontal ligament, root formation, odontoblast processes

## Abstract

**Objective:** X-linked Hypophosphatemia is associated with dental complications, including spontaneous endodontic infections (abscesses) in non-carious teeth and severe periodontal loss. Previous studies have mainly focused on dentin Hypomineralization; however, the structural basis underlying periodontal tissue failure remains unclear. We aimed to investigate histoanatomical abnormalities in the dentin and periodontium of *Hyp* mice to clarify structural consequences of *Phex* deficiency in adult molars.

**Methods:** We performed detailed histological and scanning electron microscopy analyses on the molar regions of untreated adult *Hyp* mice and wild-type littermates, with particular attention to the structural integrity of the root and periodontal ligament. Additionally, odontoblast process morphology and periodontal attachment abnormalities were evaluated.

**Results:** *Hyp* molars exhibited marked root abnormalities, including radicular shunt-like defects and disorganized odontoblast processes, particularly in furcation and radicular dentin. Periodontal attachment showed characteristic asymmetry: detachment from the cementum surface was frequently observed, whereas attachment to the alveolar bone surface was relatively preserved. These changes were accompanied by thinning and discontinuity of Sharpey’s fibers and increased vascularity in the periodontal ligament.

**Conclusions:** These findings provide a histoanatomical framework for understanding refractory dental complications in X-linked hypophosphatemia and support the importance of intervention during root development.

## 1. Introduction

X-linked Hypophosphatemia (XLH), which is caused by loss-of-function mutations in the phosphate-regulating gene with homologies to endopeptidases on the X chromosome (*PHEX*) gene, is the most common form of heritable vitamin D-resistant rickets/osteomalacia (Econs et al., 1998; Francis et al., 1995). The pathogenesis of these skeletal lesions is driven by excessive levels of fibroblast growth factor 23 (FGF23), which induce chronic Hypophosphatemia by increasing renal phosphate excretion and suppressing intestinal phosphate absorption (Haffner et al., 2025; Shimada et al., 2004). Moreover, XLH is characterized by severe oral complications, including recurrent dental abscesses without trauma or caries, as well as an increased frequency and severity of periodontitis (Foster, Nociti, & Somerman, 2014). Notably, >60% of patients with XLH suffer from periodontal defects, which result in alveolar bone loss, clinical attachment loss, and periodontal pocket formation (Arhar, Pavlič, & Hočevar, 2024; Brener, Zeitlin, Lebenthal, & Brener, 2022).

Burosumab is a fully human monoclonal antibody that targets FGF23. It effectively restores phosphate homeostasis by increasing renal phosphate reabsorption and elevating serum 1,25-dihydroxyvitamin D3 [1,25(OH)_2_D_3_] levels (Kinoshita & Fukumoto, 2018). Burosumab treatment has significantly improved skeletal phenotypes in both adults and children with XLH, as evidenced by reduced Rickets Severity Scores, alleviated joint stiffness, and accelerated healing of fractures and pseudo-fractures (Carpenter et al., 2018; Insogna et al., 2018). However, the amelioration of oral complications with burosumab treatment in patients with XLH remains insufficient (Brener et al., 2022; Seefried et al., 2023).

The *Hyp* mouse, which is an established murine homolog of human XLH, effectively recapitulates the hallmark clinical features of the disease, including chronic Hypophosphatemia and impaired bone mineralization. This model is characterized by a spontaneous 3′ deletion in the *Phex* gene; further, it has been used to investigate the molecular pathophysiology of phosphate wasting and rickets/osteomalacia (Eicher, Southard, Scriver, & Glorieux, 1976; Strom et al., 1997). Studies have investigated the dental phenotype of *Hyp* mice, with a primary focus on impaired dentin and cementum mineralization, which is often attributed to dysfunctional matrix protein production by odontoblasts and cementoblasts (Abe, Ooshima, Masatomi, Sobue, & Moriwaki, 1989; Foster et al., 2014; Zhang et al., 2020). However, despite significant advancements in systemic treatments, including conventional therapies and burosumab, there remains a persistent clinical challenge in managing oral complications in patients with XLH.

We Hypothesized that a distinct pathological mechanism, independent of Hypomineralization, may underlie the refractory dental manifestations of XLH. Accordingly, we aimed to perform a comprehensive structural analysis of the dental and periodontal tissues in 6-month-old adult *Hyp* mice. We selected this specific age to capture the terminal phenotype of the dental and periodontal attachment apparatus. Previous studies have characterized the early mineralization defects; however, the most clinically significant problems in XLH, including periodontal destruction, are often treatment-resistant and manifest as cumulative pathologies in adulthood. Moreover, recurrent abscesses occur after the eruption of deciduous teeth. Therefore, we focused on adult mice to identify the intrinsic structural defects and advanced inflammatory responses that cause these refractory oral complications. Taken together, we aimed to identify specific structural anomalies that may directly cause XLH-related oral complications in order to provide a novel anatomical basis for elucidating this treatment-resistant pathology.

## 2. Materials and methods

### 2.1. Animals

The animal experiments were conducted as per the Guidelines for Proper Conduct of Animal Experiments formulated by the Science Council of Japan. The protocols were approved by the Institutional Animal Care and Use Committee of the Osaka Women’s and Children’s Hospital: (Permit number: BMR1). All included mice were of the C57BL/6J genetic background. *Hyp* mice were initially obtained from the Jackson Laboratory (Bar Harbor, ME). Subsequently, we established six-month-old male hemizygotes (*Hyp*/Y, *Hyp*) and sibling wild-type control (*+*/Y, WT) male mice. *Hyp* genotyping was performed by genomic polymerase chain reaction (PCR) using the following primer set: *Phex* forward primer, 5′-CCAAAATTGTTCTTCAGTACACC-3′, and *Phex* reverse primer, 5′-ATCTGGCAGCACACTGGTATG-3′. A 258-bp PCR product was obtained from the WT *Phex*, but not *Hyp*, allele (Miyagawa et al., 2014). Mice were maintained in a pathogen-free barrier facility, fed standard mouse chow ad libitum, and exposed to a 12-h light/dark cycle.

### 2.2. Morphological assessment of defects in mandibular molars via scanning electron microscope (SEM)

The right mandibles and maxillae were fixed in Karnovsky’s fixative (Muto Pure Chemicals, Tokyo, Japan) at 4°C. Subsequently, samples were dehydrated through an ascending ethanol series and embedded in epoxy resin (Nisshin EM, Tokyo, Japan). Surfaces for observation were sectioned using a band saw and polished to a mirror finish using a 0.3-μm diamond abrasive sheet (Maruto, Tokyo, Japan). Backscattered electron images were acquired using a TM3000 SEM (Hitachi, Tokyo, Japan) as previously described (Takashima et al., 2024).

### 2.3. Histological evaluation of dental and periodontal tissue abnormalities in decalcified tissues

The left mandibles and maxillae were fixed in 4% paraformaldehyde for 24 h at 4°C and decalcified in 0.5 mol/l ethylenediaminetetraacetic acid. The right mandibles (decalcified for 2 weeks) were cryoprotected in sucrose and embedded in Optimal Cutting Temperature compound. Subsequently, we prepared 14-µm sagittal frozen sections using a cryostat, followed by staining using hematoxylin and eosin (H&E) or toluidine blue (pH 7.0). Next, the sections were incubated with an anti-CD31 primary antibody (bs-0468R; Bioss Inc., 1:100) overnight at 4°C, followed by incubation with an appropriate secondary antibody using a VECTASTAIN kit (Vector Laboratories). Separately, the decalcified right maxillae were embedded in paraffin, sectioned, and stained with H&E and Picrosirius Red to evaluate collagen bundle organization. All images were captured using a BZ-X1000 fluorescence microscope (Keyence, Osaka, Japan).

### 2.4. Quantitative analysis of odontoblast processes in the right mandibular first molars via phalloidin staining

To evaluate the morphological integrity of odontoblast processes (OPs) within the dentinal tubules, we performed actin cytoskeletal imaging on decalcified frozen sections of the right mandibular first molars. First, the sections were stained using Alexa Fluor 647-conjugated phalloidin (Thermo Fisher Scientific, Waltham, MA, USA) to visualize the filamentous actin (F-actin) architecture within the OPs. Fluorescence signals were captured using a fluorescence microscope. To ensure reproducibility of the analysis, OPs were imaged in three distinct anatomical dentin regions: the cusp, root furcation, and radicular sites (specifically targeting the inter-alveolar and inter-dental regions). We quantified the morphological characteristics of OPs, including their length, density, and straightness, using ImageJ software (National Institutes of Health, Bethesda, MD, USA) as previously described (Lavicky et al., 2021).

### 2.5. Statistical Analysis

Between-group comparisons were performed using Welch’s *t*-test. Data are presented as the mean ± standard error of the mean.

## 3. Results

### 3.1. Abnormal odontogenesis and dentine clefts in the crown region of *Hyp* molars

SEM identified several distinct structural defects in the mandibular molars of *Hyp* mice (Fig. 1A). *Hyp* mice showed significant dentin dysplasia and pulp chamber enlargement. The cusp region showed a multitude of longitudinal clefts throughout the dentin layer. Furthermore, the predentin layer at the pulp chamber was distinctly thickened and retained in *Hyp* mice than in WT controls. Moreover, the enamel displayed hallmark signs of Hypomineralization. Histological evaluation of the decalcified sections revealed the presence of interglobular dentin within the crown region in the *Hyp* mice (Fig. s1).

**Figure 1.**
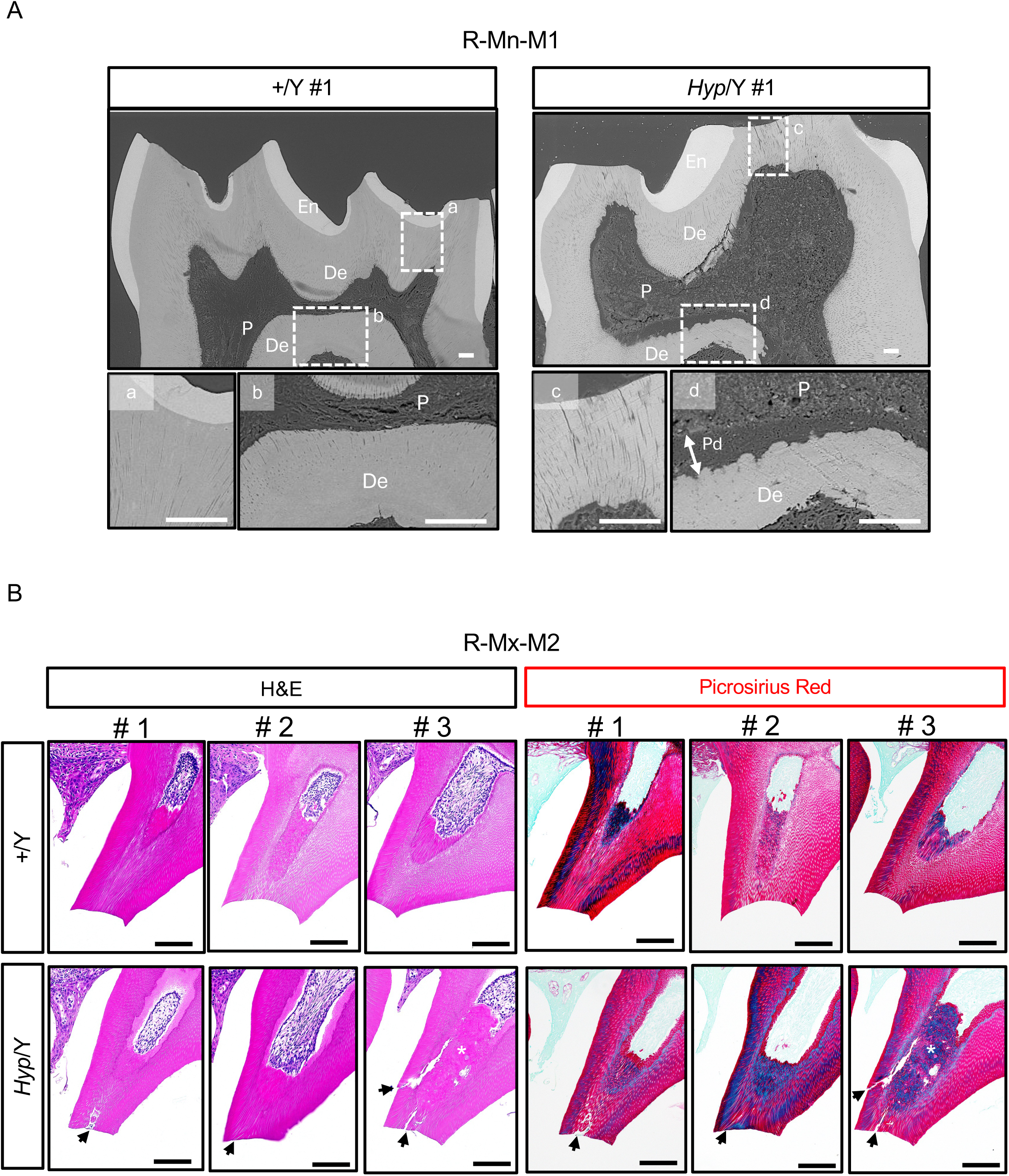
Structural alterations and dentin clefts in the crown region of *Hyp* molars. (A) Representative scanning electron microscopy (SEM) images of right mandibular first molars (R-Mn-M1) from wild-type (+/Y) and *Hyp* (*Hyp*/Y) mice. Insets (a–d) show higher-magnification views of the boxed areas. En, enamel; De, dentin; P, pulp; Pd, predentin. Scale bars, 50 μm. (B) Hematoxylin and eosin (H&E) and Picrosirius Red staining of the mesiobuccal cusp region of right maxillary second molars (R-Mx-M2). Arrows indicate longitudinal dentin clefts. Asterisks indicate reparative dentin formation. Scale bars, 100 μm.

Next, we examined decalcified paraffin sections of the maxillary molars. There was additional dentin deposition below the mesiobuccal cusps of the maxillary second molars across all genotypes (Fig. 1B). The WT group exhibited a regular tubular structure immediately adjacent to the primary dentin, which was identified as physiological secondary dentin. Contrastingly, the pattern of deposition in *Hyp* mice was dependent on the severity of structural defects. Individuals with dentinal clefts (longitudinal clefts) in the primary dentin that did not reach the pulp showed secondary dentin formation similar to that of the WT group. However, upon cleft extension to the pulp chamber, there was significant formation of reparative dentin, which exhibited a highly disrupted and irregular tissue structure.

### 3.2. Distinct structural defects in the radicular region of *Hyp* molars

Detailed SEM observations of the mandibular first molars (Mn-M1) revealed a massive shunt-like structure in the radicular of one *Hyp* mouse (Fig. 2A). Moreover, there was a corresponding anatomical feature in the radicular region of the mandible first molar (Mn-M1) in WT littermates. Contrastingly, the structure in WT mice exhibited the typical characteristics of a physiological lateral canal, with a diameter of ≈20 μm and a clear, well-defined demarcation between the dentin and cementum layers (Fig. s2A). Contrastingly, the structure observed in *Hyp* mice showed a distinctly pathological configuration. This shunt reached a maximum diameter of 50 μm and was characterized by a complete lack of normal hard tissue barriers. Notably, the pre-dentin (Pd) tissue directly extended to the periodontal ligament (PDL) space, which represented a congenital failure of odontogenesis and cementogenesis in this localized area (Fig. 2A). In the maxillary second molar (Mx-M2), there were massive shunt-like structures in the root regions of two other *Hyp* mice. Although these specific structures did not form a complete patent communication between the dental pulp and PDL, there was marked deterioration of the histological architecture of the surrounding dentin and cementum (Fig. 2B, Fig. s2B).

**Figure 2.**
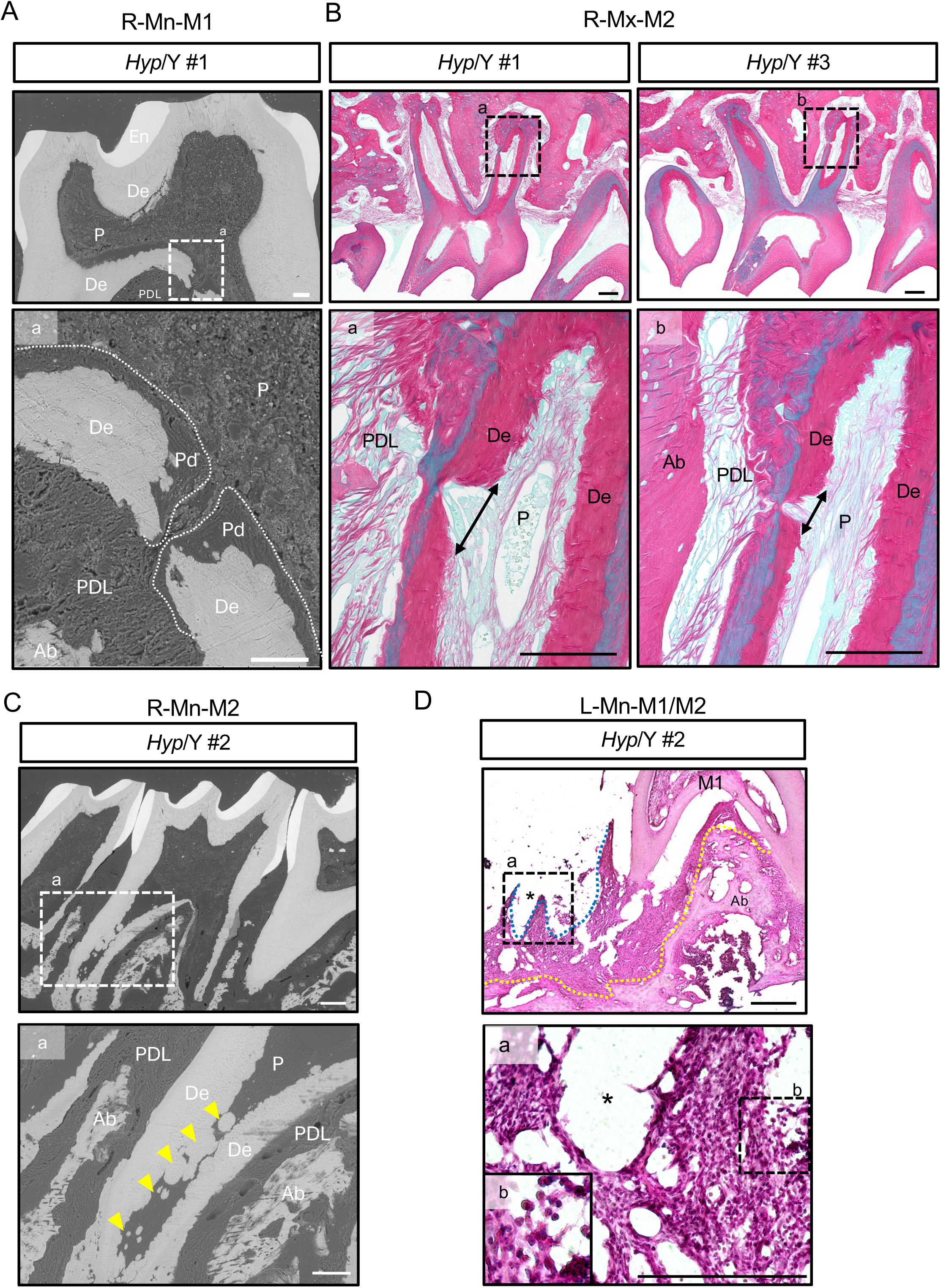
Structural abnormalities in the root region of *Hyp* molars. (A) Scanning electron microscopy (SEM) images of right mandibular first molars (R-Mn-M1) showing a shunt-like defect in *Hyp* mice. The lesion is characterized by disruption of the dentin and cementum continuity, with the predentin extending toward the periodontal ligament space (dotted line). Insets show higher-magnification views. En, enamel; De, dentin; P, pulp; Pd, predentin; PDL, periodontal ligament; Ab, alveolar bone. Scale bars, 100 μm (B) Picrosirius red-stained histological sections of right maxillary molars (R-Mx-M2) showing incomplete shunt-like defects in *Hyp* mice. Bidirectional arrows indicate the extent of the defect. (C) SEM image showing ectopic mineralization (denticle formation; yellow arrowheads) in the pulp of a *Hyp* molar. (D) Histological section of a spontaneously exfoliated molar site in the left mandibular first/second molar region (L-Mn-M1/M2) in a *Hyp* mouse, which show extensive granulation tissue and inflammatory cell infiltration replacing the normal PDL architecture. Scale bars, 100 μm.

Further, histological analysis of adult *Hyp* molars revealed pathological features extending beyond simple Hypomineralization of dental hard tissues. In the Mn-M2 radicular pulp of *Hyp* mouse #2, we identified prominent ectopic mineralization (Fig. 2C). These mineralized masses exhibited distinct dentinal tubule-like structures within their matrix; accordingly, they were identified as true denticles (pulp stones).

Finally, we confirmed spontaneous Mn-M2 exfoliation during mandibular dissection in one specific *Hyp* individual (Fig. 2D). Histological analysis of decalcified frozen sections using H&E staining revealed a prominent tissue defect corresponding to the alveolar socket of the lost Mn-M2. This region was characterized by extensive inflammatory granulation tissue with dense, diffuse inflammatory cell infiltration. The normal physiological architecture of the PDL was obliterated and replaced by this pathological tissue, which harbored a predominant population of lymphocytes.

### 3.3. Morphological analysis of odontoblast processes in mandible first molars

Accordingly, we deduced that cellular structure abnormalities, beyond mere calcification defects, might underlie the various structural anomalies observed in *Hyp* mice. Accordingly, we performed actin cytoskeletal imaging via phalloidin staining in the mandibular molars to quantitatively examine the morphology of OPs extending through the dentin of the left mandibular first molar (Mn-M1) (Fig. 3A).

**Figure 3.**
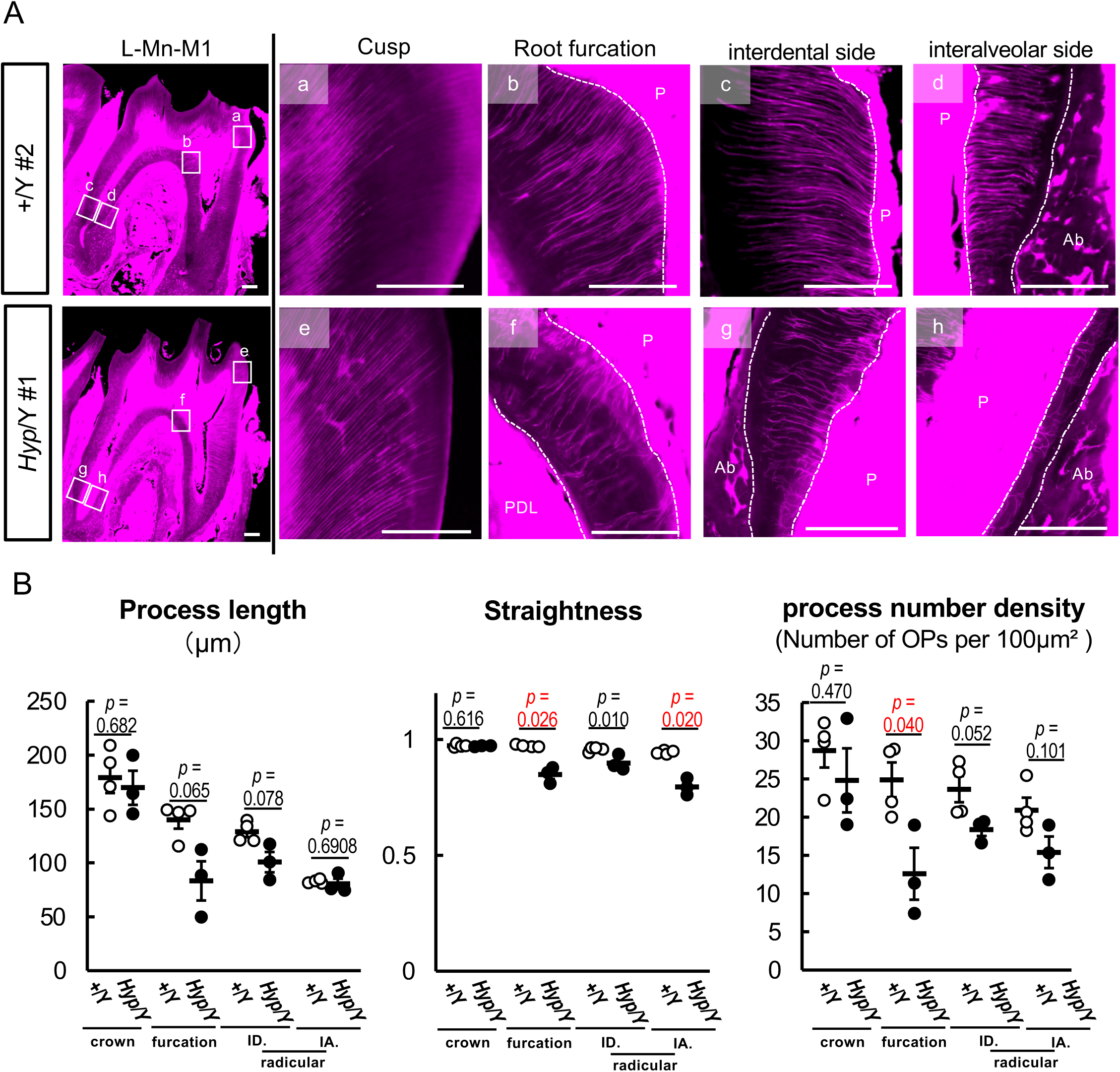
Disorganization of odontoblast processes in the root region of *Hyp* molars. (A) Phalloidin staining of odontoblast processes (OPs) in left mandibular first molars (L-Mn-M1). Representative images from crown, root furcation, and radicular regions are shown. (B) Quantitative analysis of OP morphology, including process length, straightness, and number density. Each point represents an individual measurement; bars indicate mean ± SEM. Statistical significance was determined using Welch’s *t*-test. Scale bars, 100 μm.

Quantitative analysis revealed significant morphological impairments of OPs of Mn-M1 in the region from the root furcation to the root apex in *Hyp* mice compared to their WT littermates (Fig.3B). In the cusp region of dentin, there were no significant between-group differences in the length, straightness, or density of OPs. Contrastingly, the radicular dentin in *Hyp* mice, which was the root furcation and root regions, exhibited significant deterioration in the straightness or density of OPs compared to WT mice. Notably, OPs in the root furcation of *Hyp* mice showed the most pronounced structural impairment, with a significant decrease in both straightness and density compared with WT littermates (*p* < 0.05).

### 3.4. Evaluation of periodontal ligaments (PDLs) in maxillary molars

Given the spontaneous tooth loss associated with severe periodontitis in the mandibular molar of a *Hyp* mouse, we evaluated the structural integrity of PDLs in the right maxillary second molars (Mx-M2) using histological observation (Fig. 4A and Fig. s3A).

**Figure 4.**
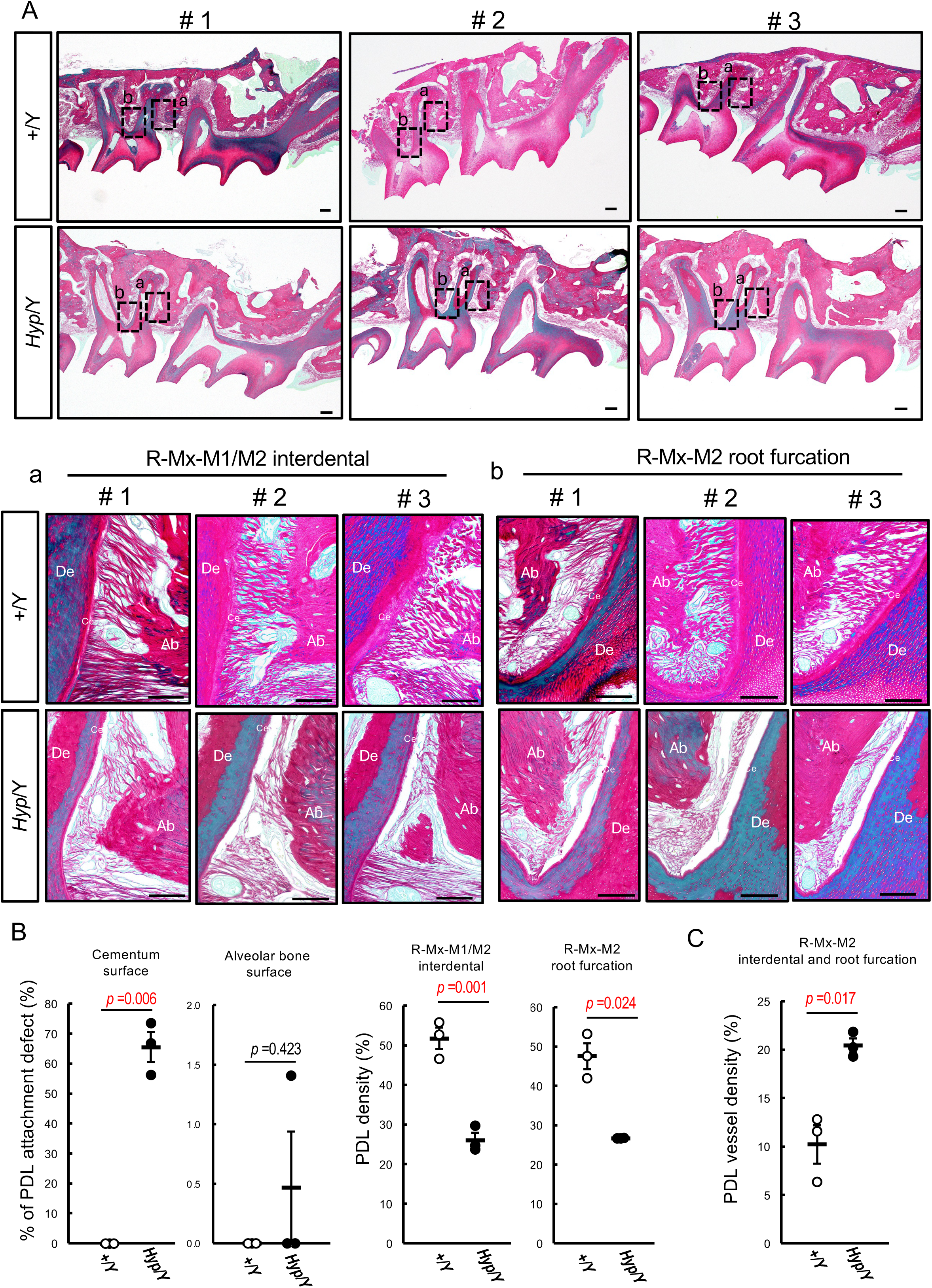
Impairment of periodontal attachment and increased vascularity in *Hyp* mice. (A) Picrosirius Red-stained sections of right maxillary molars showing the interdental region between the right maxillary first and second molars (R-Mx-M1/M2; a) and the root furcation region of the right maxillary second molar (R-Mx-M2; b). *Hyp* mice show thinning and discontinuity of periodontal fibers, with frequent detachment of the periodontal ligament (PDL) from the cementum surface. De, dentin; P, pulp; Ce, cementum; Ab, alveolar bone. Scale bars, 100 μm. (B) Quantification of PDL attachment defects at the cementum and alveolar bone surfaces. (C) Quantification of PDL vessel density, showing increased vascularity in *Hyp* mice. Each point represents an individual sample; bars indicate mean ± SEM. Statistical significance was determined using Welch’s *t*-test.

In the Picrosirius Red-stained sections, the Sharpey’s fibers within the Mx-M1/M2 interdental and Mx-M2 furcation regions were markedly thinner and more discontinuous in *Hyp* mice than in WT controls. Further, in *Hyp* mice, although transseptal fibers connecting adjacent teeth maintained relatively strong connectivity, there was pathological PDL detachment from the cementum surface (Fig. 4A and Fig. s3A). Quantitative analysis revealed that the PDL attachment defect area at the cementum interface averaged 65% in *Hyp* mice, whereas there was no such detachment in WT littermates. Contrastingly, the PDL insertion at the alveolar bone surface remained intact in both groups (Fig. 4B).

In addition to the structural detachment, we evaluated the vascularity of the periodontal tissues. Histological observations indicated that the high vascularization of PDL in *Hyp* mice. Specifically, the blood vessel density in the maxillary M2 PDL was approximately two-fold higher in *Hyp* mice than in WT littermates (Fig. 4C). Furthermore, this Hyper-vascularization was confirmed in the mandibular M1 and M2 regions (Fig. s3B). Moreover, the PDL of *Hyp* mice exhibited dense accumulation of CD31-positive cells, which is a marker for vascular endothelial and immune cells (Fig. s3C).

## Discussion

Our findings provide new histoanatomical evidence that oral complications of XLH cannot be solely explained by Hypomineralization of dental hard tissues; instead, they are characterized by profound architectural failure at the root-periodontium interface. Adult *Hyp* mice showed marked radicular abnormalities, including shunt-like defects (Fig. 2A, 2B), disorganization of odontoblast processes (Fig. 3), and selective PDL detachment from the cementum surface (Fig. 4A, 4B). These structural defects were accompanied by Hypervascular and inflammatory changes in the periodontal tissues (Fig 2D, 4C, s3B, s3C). Taken together, our findings suggest that the refractory oral manifestations of XLH arise from congenital and developmentally established structural vulnerabilities that compromise the integrity of the dentin-periodontium complex.

The crown phenotype observed in this study was broadly consistent with previous descriptions of XLH and *Hyp* mice(Arhar et al., 2024; Chaussain-Miller et al., 2007; Opsahl Vital et al., 2012), including enlarged pulp chambers, dentin dysplasia, interglobular dentin, and enamel Hypomineralization (Fig. 1A, s1). These abnormalities have traditionally been interpreted as consequences of impaired mineralization, which leads to increased susceptibility to crack formation, bacterial invasion, and recurrent abscesses. However, our findings suggest that these coronal changes do not solely account for the entire spectrum of oral pathology in XLH. Several *Hyp* teeth showed crown structural defects without overt pulp destruction, with secondary or reparative dentin formation remaining detectable beneath the cusps (Fig 1B). This indicates that, despite the abnormal dentin matrix environment, some defensive or compensatory capacity of the pulp may be retained.

Contrastingly, the root region showed different structural abnormalities. Notably, there were distinctly large shunt-like lesions in the radicular region relative to the small physiological lateral canals occasionally observed in WT mice (Fig 2A, 2B, s2). These lesions showed disrupted continuity of the dentin; further, there was abnormal predentin extension toward the PDL space (Fig 2A). This suggests a localized failure of coordinated root hard tissue formation rather than a simple quantitative reduction in mineral deposition. Given the pivotal role of the root surface and its periodontal attachment apparatus in maintaining long-term tooth stability, such developmental defects may represent a key anatomical substrate for the progressive and treatment-resistant dental complications observed in XLH.

Our analysis of OPs further highlights the vulnerability of root development in *Hyp* mice (Fig 3). In the crown, the OP morphology was relatively preserved, while they showed reduced straightness and density in the furcation and radicular regions. This regional difference is important given that root dentin is formed during a dynamic post-eruptive phase requiring highly coordinated odontoblast activity. Therefore, our findings suggest non-uniform architectural disorganization in *Hyp* teeth throughout odontogenesis, which becomes most pronounced during root formation. This interpretation is consistent with clinical observations indicating that early treatment in patients with XLH may attenuate subsequent dental complications, especially when initiated prior to the completion of permanent tooth development (Kato et al., 2022). Our findings support the concept that preserving phosphate homeostasis during the critical window of root formation is crucial for establishing proper dentin architecture.

Another major finding was the highly selective nature of periodontal attachment failure. In *Hyp* mice, the PDL frequently detached from the cementum surface, whereas its insertion into the alveolar bone remained relatively preserved (Fig 4A, 4B, s3). This asymmetry is notable since it indicates that the cementum-side attachment apparatus may be vulnerable in XLH. Sharpey’s fibers were thinner and more discontinuous in *Hyp* mice, which further supports the notion that the cementum-PDL interface is intrinsically compromised (Coyac et al., 2017; Phanrungsuwan et al., 2025). Such a defect may have substantial pathological consequences; specifically, upon weakening of the root-side attachment, the tooth becomes chronically susceptible to microbial ingress and inflammatory extension into deeper periodontal tissues, even with initial intactness of the alveolar bone-side attachment.

This structural vulnerability was accompanied by clear evidence of an altered periodontal microenvironment. In *Hyp* mice, the PDL showed increased vascularity and accumulation of CD31-positive cells (Fig. 4C, s3B, s3C), which is consistent with an inflammatory and Hyper-reactive tissue state. These findings suggest that the periodontal tissues in XLH are not merely passively weakened but rather may exist in a condition of chronic predisposition to inflammation. Accordingly, structural defects such as radicular breaches and cementum detachment may facilitate exposure to external stimuli and, therefore, promote persistent tissue activation. Osteopontin accumulation has been implicated in XLH-associated cementum defects and periodontal pathology; moreover, local dysregulation of mineralization and immune responses may synergistically exacerbate periodontal breakdown (Boukpessi et al., 2017; Phanrungsuwan et al., 2025). Although we did not directly assess this mechanism, our histological findings are compatible with such a model.

Notably, one *Hyp* mouse showed spontaneous tooth exfoliation (Fig. 2D). Tooth loss at this age is highly atypical in WT mice (An et al., 2017; Liang, Hosur, Domon, & Hajishengallis, 2010), suggesting that the loss of periodontal integrity in *Hyp* mice occurs prematurely and pathologically rather than as part of physiological aging. Histological examination of the affected site showed extensive inflammatory granulation tissue and complete disruption of normal periodontal architecture. This supports the notion that tooth exfoliation in XLH does not simply result from weak mineralized tissues, but rather a destructive process driven by the interaction of congenital structural defects and secondary inflammatory breakdown. Specifically, the functional lifespan of the periodontium may show early shortening due to incomplete and abnormal construction of the root attachment apparatus.

Notably, we focused on untreated adult *Hyp* mice. Previous studies have extensively characterized early mineralization defects in developing teeth (Coyac et al., 2017; Ogawa et al., 2006; Zhang et al., 2020); however, fewer studies have examined the terminal structural consequences that become apparent after long-term function. Analysis of adult animals allowed identification of pathological features possibly more directly relevant to the observed refractory oral complications, including periodontal collapse and spontaneous tooth loss. Accordingly, the *Hyp* mouse remains pivotal for assessing phosphate metabolism as well as how developmental defects in the dentin-periodontium complex translate into cumulative structural failure in adulthood.

This study has several limitations. First, we included a limited number of animals, and some lesions were not uniformly observed in all teeth or individuals. Second, the study was restricted to a single age point, which impeded temporal analysis of the emergence of each abnormality. Third, although our findings strongly indicate structural and inflammatory abnormalities, the molecular mechanisms linking Phex deficiency to these regional defects remain unclear. Future studies combining longitudinal analysis with molecular and therapeutic approaches are warranted to determine how phosphate dysregulation, local matrix abnormalities, and periodontal inflammation interact during disease progression.

## Conclusion

Our findings expand the current understanding of XLH oral pathology by identifying root-periodontium architectural failure as a central component of disease pathogenesis. Beyond the established concept of dentin and enamel Hypomineralization, XLH appears to involve developmental defects in root construction and periodontal attachment that lead to long-lasting predisposition to inflammatory breakdown. Our findings provide an anatomical framework for understanding the severity and refractoriness of oral complications in XLH.

## Author Contributions

**Chiaki Nishizawa:** conceptualization, data curation, formal analysis, investigation, methodology, visualization, writing – original draft, writing – review and editing, funding acquisition. **Jiro Miura:** methodology, resources, investigation, data curation, writing – review and editing. **Tomoaki Iwayama:** Validation, writing – review and editing. **Miwa Yamazaki:** Resources, writing, review, and editing. **Toshimi Michigami:** validation, writing – review and editing, supervision. **Kazuaki Miyagawa:** project administration, supervision, funding acquisition, conceptualization, data curation, formal analysis, investigation, methodology, visualization, writing – original draft, writing – review and editing.

## Acknowledgments

We extend our sincere gratitude to Dr. Yuki Matsushita (Graduate School of Biomedical Sciences, Nagasaki University) and Dr. Mizuki Nagata (Graduate School of Medical and Dental Sciences, Institute of Science, Tokyo) for their valuable advice and technical guidance on the preparation of frozen sections.

We thank Editage (http://www.editage.jp) for the English language editing.

## Ethics Statement

All animal experiments were performed in accordance with the guidelines of the Institutional Animal Care and Use Committee of the Osaka Women’s and Children’s Hospital (Permit number: BMR1).

## Data Availability Statement

The authors confirm that the data supporting the findings of this study are available within the article.

## Declaration of Generative AI and AI-assisted technologies in the writing process

During the preparation of this manuscript, the authors used ChatGPT (OpenAI, San Francisco, CA, USA) to assist with language refinement and improve clarity. All content was subsequently reviewed and edited by the authors, who take full responsibility for the integrity and accuracy of the work.

