## Supplemental figures for "Radicular and periodontal structural defects underlie refractory oral pathology in the adult *Hyp* mouse model of X-linked Hypophosphatemia"

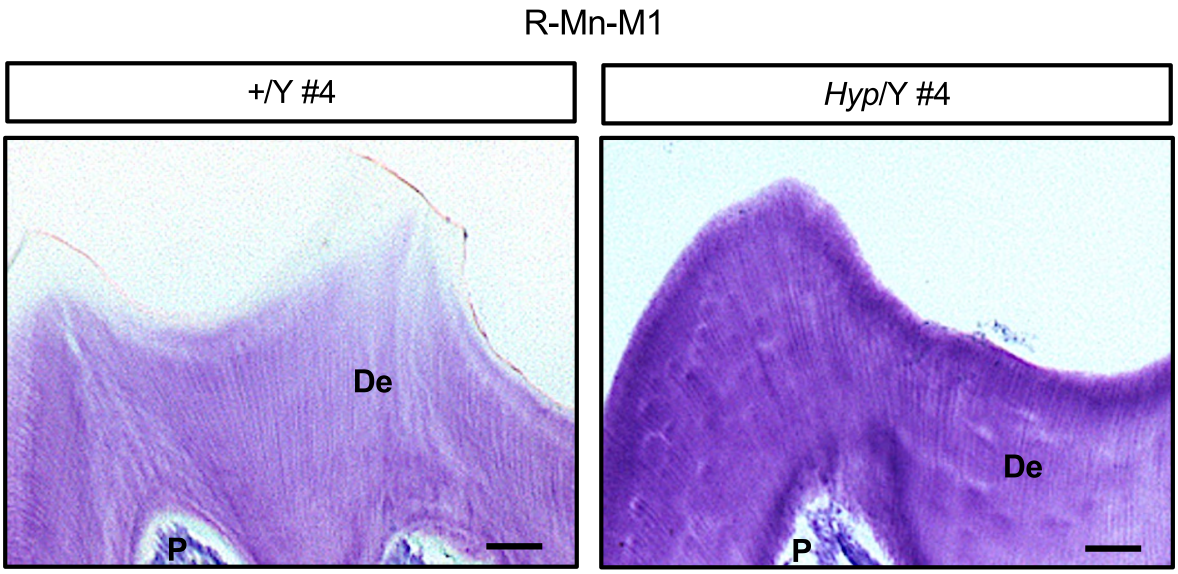


**Figure S1. Structural alterations in the crown region of *Hyp* molars (additional examples).** Representative histological images of right mandibular first molars (R-Mn-M1) from wild-type (+/Y) and *Hyp* (*Hyp*/Y) mice. Toluidine blue staining show dentin dysplasia, enlarged pulp chambers, and irregular dentin structure in *Hyp* mice. De, dentin; P, pulp. Scale bars, 50 μm.

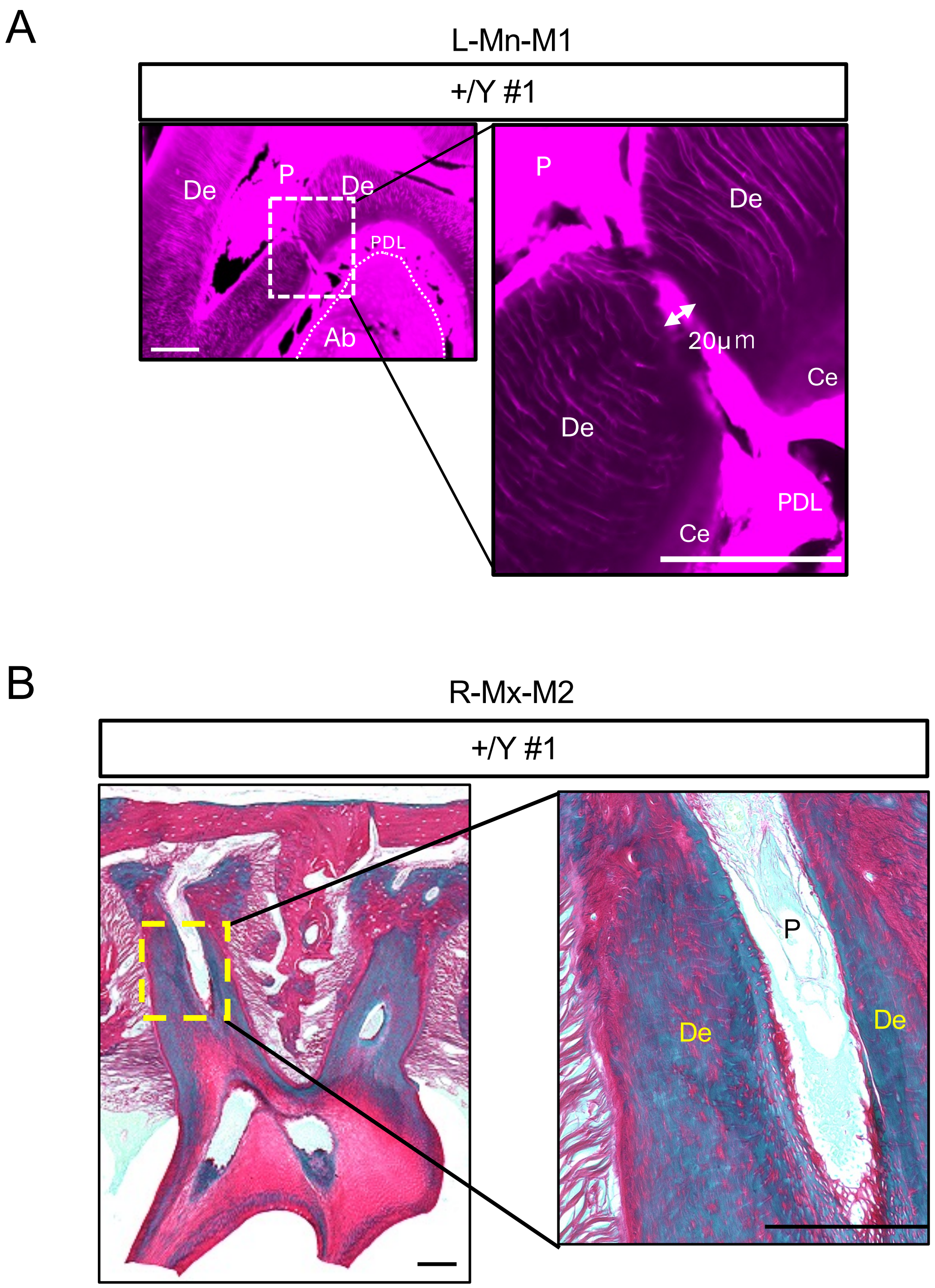


**Figure S2. Physiological structures in wild-type molars.**

(A) Representative phalloidin-stained image of a left mandibular first molar (L-Mn-M1) from a wild-type (+/Y) mouse. A physiological lateral canal (arrow) is observed, showing a narrow and well-defined communication between the pulp and periodontal ligament (PDL), with preserved dentin–cementum continuity. (B) Representative Picrosirius Red-stained image of a right maxillary second molar (R-Mx-M2) from a wild-type (+/Y) mouse. The inset shows a higher-magnification view of the boxed area. De, dentin; P, pulp; Ce, cementum; Ab, alveolar bone; PDL, periodontal ligament. Scale bars, 100 μm.

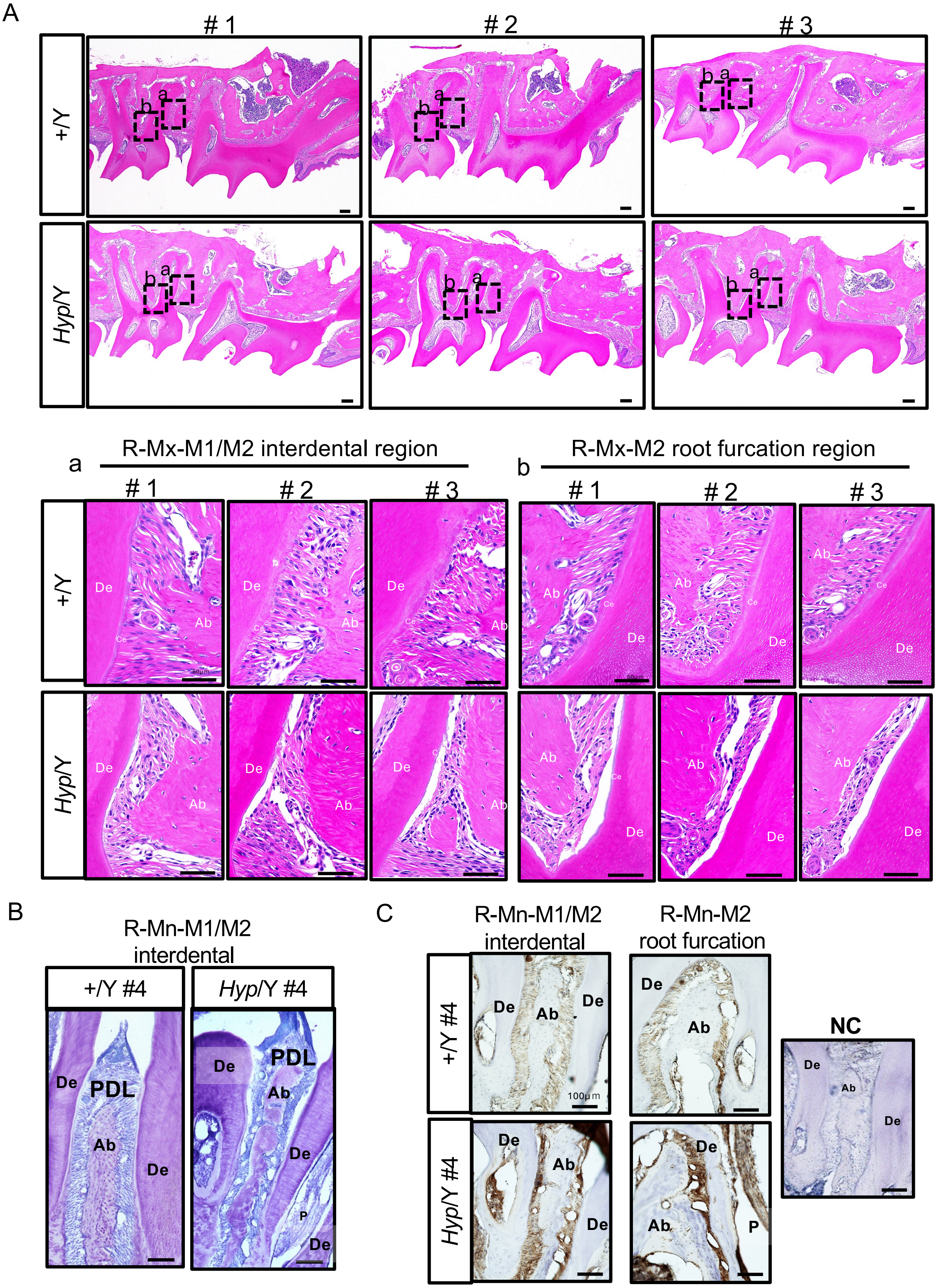


**Figure S3. Additional evidence of periodontal attachment defects and increased vascularity in *Hyp* mice.** (A) H&E-stained sections of right maxillary molars (R-Mx-M1/M2) from wild-type (+/Y) and *Hyp* (*Hyp*/Y) mice (#1–#3). (B) Toluidine blue-stained higher-magnification views showing detachment of the periodontal ligament (PDL) from the cementum surface of right mandibular molars (R-Mn-M1/M2) in *Hyp* mice. (C) CD31 immunostaining showing increased vascularity of right mandibular molars (R-Mn-M1/M2, R-Mn-M2) in the PDL of *Hyp* mice. De, dentin; Ab, alveolar bone; Ce, cementum; PDL, periodontal ligament. Scale bars, 100 µm.
